# Exploring dog-owner microbiota and bacterial pathogens in Southeast Asia

**DOI:** 10.64898/2026.10.07.757238

**Authors:** Pauline Van Leeuwen, Anamika Kritiyakan, Serge Morand, Areeya Kriengudom, Alix Nicolas, Michel de Garine Wichatitsky, Johan Michaux

**Author notes:** **Corresponding author:** Johan Michaux.

## Abstract

Domestic dogs may influence human microbiota and act as reservoirs or sentinels for zoonotic bacteria, yet little is known about these dynamics in Southeast Asia. Using full-length 16S rRNA gene sequencing from rectal swabs, we analyzed microbiota from dog-owner pairs from four villages of rural Thailand (Nan province) to assess microbial diversity, shared taxa, and zoonotic potential. Dogs exhibited higher microbial and putative pathogen diversity than humans, particularly during the rainy season. While dog-owner pairs shared more bacteria than unrelated individuals, overall overlap remained limited — likely driven by their dogs’ free-roaming lifestyles and broader environmental exposures. However, co-occurrence of putative zoonotic pathogens such as *Shigella* spp. and *Klebsiella pneumoniae* suggests dogs could serve as useful indicators for specific public health risks, with further targeted pathogen screening needed to validate these results. Our findings highlight the value of integrating dogs into zoonotic surveillance programs, particularly in regions with free-ranging dog populations and close human-animal contact.

## 1. Introduction

The human microbiota is a cornerstone of health, influencing key physiological and immunological processes that determine metabolism, disease susceptibility and the capacity to respond to infections. Among the myriad factors shaping human microbial communities, interactions with domestic animals — especially companion dogs — may play an underestimated role. The dog was the first domesticated species by humans, interacting as a commensal in human societies for more than 30,000 years (Savolainen et al., 2022). Domestic dogs provide multiple (socio-economic, cultural, recreational, mental and physiological health) benefits (Gray & Young, 2011), while also negatively impacting the health of humans (zoonosis, Stull et al., 2015), and of the environment including wildlife (Hughes & Macdonald, 2013). Dogs share human environments, sometimes including their homes, and even their behaviors, creating ample opportunities for microbial exchange.

People may acquire pet-associated zoonotic infections through bites, scratches or other direct contact of their skins or mucous membranes, direct contact with animal saliva, urine and other body fluids or secretions, ingestion of animal fecal material, inhalation of infectious aerosols or droplets and through the bite of arthropods and other invertebrate vectors. While several zoonotic pathogens (*e.g.*, pathogenic *Escherichia coli*, *Salmonella sp., Campylobacter sp., Brucella canis*) have been identified in dogs, their potential as bacteriome proxies for humans remains underexplored (Stull et al., 2015; Ghasemzadeh & Namazi, 2015). The overall microbiota of dogs, like that of humans, is dynamic and influenced by intrinsic factors such as breed and health conditions (Deschamps et al., 2022; Suchodolski et al., 2012) and extrinsic factors including diet and environment (Kim et al., 2017; Zhao et al., 2023). As shown by Yarlagadda et al. (2022), dogs in Laos had greater gut microbial diversity compared to dogs from the USA, South Africa and India, which was not solely explained by diet. Moreover, social behaviors, such as grooming and proximity to humans and other dogs, further contribute to microbiota composition (Coelho et al., 2018). Despite this complexity, dogs have been found to exhibit greater microbial diversity than humans (Wetzels et al., 2021), and to share more gut microbes with humans than other mammals (Coelho et al., 2018), suggesting a potential role for the monitoring of zoonotic disease risks. Similar to humans, domestic dogs have an omnivorous metabolism and can digest, absorb and metabolize dietary carbohydrates, illustrating gut microbial similarity to humans, as compared to mice or pigs. However, a recent study conducted in Hungary showed that humans and pet dogs exhibit less similar skin bacterial communities compared to humans and wolves, whereas all canines tested appeared grouped more closely together, leaving humans as a clear separate cluster (Wetzels et al., 2021) and suggesting greater importance of host species than host environment.

Studies investigating dog-human microbiota shown two main recurrent results: (1) Owners in households with dogs exhibit greater microbial diversity (, based on gut/fecal samples (Kates et al., 2020), household environmental surface swabs (Dunn et al., 2013), and paired human-pet skin swabs at Staphylococcus carriage sites (nares, inguinal/axilla; Misic et al., 2015); (2) Owners have more similar microbiota to their dogs than to humans outside of their household, based on paired fecal samples (Ito et al., 2024), fecal Enterococcus isolates (Damborg et al., 2009), and skin, oral, and fecal samples from cohabiting families (Song et al., 2013).. These studies have focused on Europe and North America, leaving a critical gap in understanding the dynamics of dog microbiota in other parts of the world, including Southeast Asia, where pet ownership practices and environmental contexts differ markedly (Hernandez et al., 2022). In Western and/or industrialized countries, dogs are kept mostly for pleasure and company and are often considered family members or friends (Blouin, 2013; Gillet et al., 2024; Kim et al., 2020; Smith et al., 2022). On the other hand, in low-and middle-income countries, the majority of dogs have a free-ranging lifestyle and are kept primarily for guarding purposes, followed by companionship (Massei et al., 2017; Marshall et al., 2023; Sambo et al., 2014). In rural Southeast Asia, free-roaming dogs are owned by a single owner or by a community and are not fully controlled in terms for their movements and health management. Recent bibliometric analysis underscores the importance of considering canine ecotypes — defined by their ecological roles and living conditions — in zoonotic transmission dynamics (Langlois et al., 2025). Compared to pet dogs, free-roaming dogs have more mobility, exposure to contaminated environments, and interactions with multispecies communities that make them important contributors to pathogen adaptation and spillover. Thus, frequent contact between free-ranging dogs and humans makes them potential vectors for the direct transmission of pathogens and contributors to the spread of zoonotic agents across species. This highlights the need for ecotype-specific research and intervention strategies. Recognizing the distinct epidemiological roles of different dog populations will be critical for integrating canine data into global One Health strategies aimed at preventing pathogen spillover and enhancing disease surveillance and control.

Addressing the knowledge gap regarding dogs’ epidemiological importance for zoonotic diseases in Southeast Asia (Indonesia, Cambodia, and Thailand), SEAdogSEA project activities combine experimental approaches, social surveys and modelling to explore several aspects of dog-human interactions. This component of the project focuses on the microbiota of dog-owner pairs from rural Thailand focusing on microbial diversity, shared taxa, and pathogen profiles to better assess the potential role of dogs as a “bridge species” for the transmission of emerging zoonoses and their potential as a “sentinel”, for improved zoonotic diseases surveillance and management. We hypothesized that dogs, due to their intricate relationship with humans, could serve as proxies for the prevalence of zoonotic bacteria in their respective owners.

This study employs advanced metabarcoding sequencing of the full 16S rRNA gene to analyze microbiota composition in dog-owner pairs from Nan Province, Thailand. Compared to other metabarcoding studies that could only speculate about bacterial transmission at the genus level (Liyanagama et al., 2023), we used long reads of third generation sequencing to obtain species-level affiliation. Despite inherent bias of metabarcoding using a 16S marker, long reads allow better taxonomic assignment than short reads, although many species complexes cannot be distinguished and pathogenicity can only be putative based on literature review at the species level. By assessing microbial overlap and potential transmission routes, our objectives were to (1) characterize the human putative pathogen load in domestic dogs in rural Southeast Asia, (2) establish whether dogs are closer to their owners than to other dogs or humans and (3) assess the applicability to consider dogs as proxies for human microbial health, contributing to public health strategies and One Health initiatives.

## 2. Methods

### 2.1 Sample collection

A total of 96 rectal swabs samples (stored in 70% ethanol) from twenty-four dog-owner pairs were collected in January and July 2023 at four adjacent rural villages from Nan province (Thailand). January and July sampling timepoints correspond respectively to the dry season (November-March, with minimal rainfall) and the rainy season (April/May-October, peaking in July-August) in Nan Province, northern Thailand. We selected twenty-four adult dog-owners from four villages which participated in SEAdogSEA project surveys, including dog GPS tracking, owner questionnaire survey, and dog and dog owner samples (blood and feces). Villages sampled were Ban Huak (19°08’14.7“N, 100°45’36.5”E), Ban Nam Krai (19°07’54.5“N, 100°44’19.7”E), Ban Huay Muang (1 9°08’21.3“N, 100°43’09.3”E) and Ban Santisuk (19°07’46.2“N, 100°41’58.8”E). Each pair (an adult owner and their dog) was sampled twice in January and July 2023 using rectal swabs. One owner did not participate in the July session. Dogs were physically restrained by their owners while the certified, experienced veterinary staff collected the rectal swabs. The human samples were collected by certified staff of the local Primary Care Health Unit (PCHU). The swabs were preserved in 70% ethanol immediately after rectal swabbing in the field (dog) or at PCHU laboratory after transfer in refrigerated conditions, before further analysis at the laboratories (details in 2.2).

Informed consent was obtained verbally from each participant by local research assistants attached to the project, and witnessed by the staff from the local PCU who proceeded with sample collection and processing. The volunteer participants were reminded in Thai language of the objectives of the project and the confidentiality associated with the personal data provided and the results of the analysis that will be used for research purposes only. Volunteers approved all sample collections from their dog and themselves, after being explained in detail the overall objectives of the study and the details of procedure. The human sampling procedure was approved by the ethical committee of the Department of Health of Nan Province (permit number NAN REC 64-13), and the animal procedure was approved by ethical committee of Kasetsart University Animal Ethics (permit number ACKU65-VTN-001).

### 2.2 DNA extraction and sequencing

Using the QIAamp Fast DNA Stool Mini Kit from Qiagen, DNA from the rectal swabs was extracted following the manufacturer’s protocol, treating swabs as fecal material. An extraction blank was added to each extraction batch (n=9). The approach used for this study was metabarcoding of the full 16S rRNA gene for bacterial profiling, through a two-step PCR strategy. In PCR_1_, we amplified the full 16S rRNA gene using one primer pair (27F-1492R, Frank et al., 2008). PCR_1_ reactions were carried out using Q5 HiFi NEB polymerase, in a final volume of 25 μL containing 0.25 μL of 0.02 U/μL Q5 High-Fidelity DNA polymerase, 0.5 μL of 10 mM dNTPs, 1.25 μL of each of the 0.5 μM forward and reverse primers, 5 μL of 5x Q5 buffer, 5 μL of DNA extract and additional molecular grade water. The thermocycler optimized conditions included an initial denaturation at 98°C for 1 min, followed by 30 cycles of denaturation at 98°C for 20 s, hybridization at 61°C for 30 s and extension at 72°C for 2 min, and a final extension at 72°C for 2 min. PCR_1_ negative controls were included in every 96-well plate reaction (n=2). In PCR_2_, to trace down the sample ID in downstream analyses, we incorporated a sample-specific combination of 60 bp long in-house tags for Ligation Sequencing Kit (SQK-LSK11 2). PCR_2_ was carried out in a final volume of 25 μL, containing 0.25 μL of 0.02 U/μL Q5 High-Fidelity DNA polymerase, 0.5 μL of 10 mM dNTPs, 2.5 μL of each of the 0.5 μM forward and reverse primers, 5 μL of 5x Q5 buffer, 5 μL of PCR_1_ product and additional molecular grade water. The thermocycler optimized conditions were identical to the PCR_1_ steps except that only 7 cycles were performed with hybridization at 62°C for 30 s. All pre-PCR mixes were set up under a sterile hood in a DNA-free room to minimize the risk of contamination. Following DNA purification, quantification and pooling at equimolarity, DNA library construction and sequencing were conducted at the University of Liège GIGA Genomics platform. Finally, a total of 96 samples, 9 extraction blanks and 2 PCR_1_ blanks were sequenced on an a GridION (Oxford Nanopore Technologies) at 100,000 mean sequencing depth per sample with parameters and protocols for optimal coverage and accuracy.

### 2.3 Bioinformatics and statistical analysis

After basecalling and barcode demultiplexing using *Guppy*, we controlled the quality of these demultiplexed reads with the *FastQC* software (Andrews, 2010) and used *cutadapt* (Martin, 2011) to trim primers for all sequence reads. The *RESCUE* pipeline for cleaning and clustering of raw sequence data was used, with the RDP database for taxonomic assignments at the species level (Petrone et al., 2023). The *decontam* package in R (R Core Team, 2022) was used to detect true contaminants present in extraction and PCR_1_ blanks, resulting in 31 true bacterial contaminants that were removed from the dataset. Blank samples were removed, and taxa not belonging to the kingdom Bacteria were also removed, resulting in a total of 2176 bacteria taxa. Finally, due to great variation in sampling depth between samples, we rarefied data to 20,000 reads per samples based on rarefaction curve (Supp Figure 1), removing samples with less than 20,000 reads resulting in a final dataset of 87 samples (Supp Table 1) and 2124 bacterial taxa for alpha diversity analysis.

Tests described below were conducted on the human and dog microbiota following separation of bacterial taxa based on their putative pathogenic status for humans. First, we retrieved taxa assigned as established pathogens for humans from the database established by Bartlett et al. (2022). Their transmission route was assessed through a review of the literature (Bergey’s Manual 2001-2012). Additionally, human pathogenic bacterial species for which dogs can be competent hosts were documented with the existing literature (Langlois et al., 2025). Other taxa without established pathogenicity for humans were categorized as symbionts. Each category was considered mutually exclusive. The proportion of pathogenic bacterial species is overestimated, as our methodology does not allow strain-level identification. For instance, *E.coli* is categorized as a pathogenic for humans, although many strains are known commensals of the mammalian flora. Alpha diversity (within-sample diversity) was quantified using the observed number of phylotypes (lowest taxonomic level) and Shannon index (estimated community diversity) and was investigated using linear models. Alpha diversity for symbionts, human pathogens, and pathogens where dogs are competent was modeled using linear mixed-effects models with host species, village, and sampling date as fixed effects, and individual identity nested within dog-owner pair as a random effect, to account for repeated sampling and pair-level non-independence. The estimates associated with the effect of each covariate in these models were tested using ANOVAs, after verification of model residuals that met the assumptions of normality and of homoscedasticity. Non-parametric tests (Kruskal–Wallis [KW] followed by Dunn post hoc tests when necessary) with Benjamini-Hochberg correction were used to investigate the proportion in shared taxa within dog-owner pairs and unpaired dog and humans.

Beta diversity was calculated as the Bray-Curtis distance and was visualized using principal coordinates analysis (PCoA), built using the R package vegan (Oksanen et al., 2017). Permutational multivariate analyses of variance (PERMANOVAs) implemented using the adonis function (n = 9999 permutations) were used to test similar models as for the alpha diversity, constrained within individual identity. Thereby, the effect of host species on the similarity of bacterial communities was tested among symbionts and putative human pathogens. Differences in within-group variation were investigated using a betadisper test. Bacterial community resemblance through Bray-Curtis distances within dog-owner pairs and between dog and non-owner pairs were tested with non-parametric tests (KW followed by Dunn post hoc tests when necessary) with Benjamini-Hochberg correction.

A linear discriminant analysis (LDA) was used to assess the abundance of putative pathogens between humans and dogs based on their transmission routes with a threshold logarithmic LDA score set at 3.0 and ranked, with p-value threshold at 0.05. Finally, to reveal changes in microbe community structure between dogs and humans, we used covariation network analysis as implemented in NetCoMi R package (Peschel et al., 2020) using the SPRING method. The SPRING (Semi-Parametric Rank-based approach for INference in Graphical models) method is used to construct microbial association networks by estimating sparse, rank-based partial correlations, which are robust to non-normal data distributions and compositional effects common in microbiome datasets. This network analysis was used in an exploratory capacity to visualize potential community-level co-occurrence patterns, rather than to test specific statistical hypotheses. Features present in less than 20% of the samples were removed before analysis. We estimated networks for dog and human hosts at the genus level (abundance filter threshold = 0.005; mb method; greedy clustering).

## 3. Results

### 3.1 Dogs exhibit greater microbial diversity, with more shared taxa between dogs

Overall, 2124 bacterial taxa were detected, composed of 1947 symbionts and 272 established human pathogens including 53 species for which dogs are competent hosts. Overall, dogs harbored a greater number of taxa for symbionts (n_symbiont_ = 1419) and pathogens (n_pathogen_ = 207, n_dog-competent_ = 49 where dog competent) compared to humans (n_symbiont_ = 820, n_pathogen_ = 127, n_dog-competent_ = 18). Proteobacteria and Firmicutes dominated across all samples, and dogs carried a higher relative abundance of Tenericutes compared to humans (Supp Figure 2A). For symbionts, greater overall species richness was observed in dogs despite not reaching significance, while humans exhibited greater evenness in their species richness (Observed number of taxa host effect: F = 14.66, df = 1, 38.8, p <.001; Shannon ANOVA: F = 26.4482, df = *1,* 21.04, p <.001; Figure 1 A). Dogs carried on average a slightly lower proportion of putative pathogens in their gut microbiome (12.7% ± 1 %) compared to humans (14.4% ± 2%, Supp Figure 2B). However, dogs exhibited greater species richness for putative pathogens than their owners, with a notable increase in the rainy season (Figure 1B&C), regardless of their location (linear mixed model for Observed number of taxa: host effect, F = 10.08, df = 1, 21.8, p =.004; date collection effect, F = 10.57, df = 1, 38.9, p =.002; village effect, F = 0.65, df = 3, 15.6, p =.59). Post hoc contrasts confirmed a significant increase in richness from January to July in both dogs (estimate = 5.23 ± 1.61, t = 3.24, p =.002) and humans (same contrast, p =.002). Similar results were obtained for putative pathogens for which dogs are competent hosts (host effect: F = 167.83, df = 1, 21.6, p <.001, date collection effect: F = 15.20, df = 1, 39.0, p <.001). Food and oro-fecal transmission routes predominated in dogs for putative bacterial pathogens (22.1% ± 3% and 25.6% ± 3% respectively, Supp Figure 2C). Oro-fecal transmission was significantly more abundant in dogs than in their owners (linear mixed model with individual identity nested within dog-owner pair as a random effect, transmission × host interaction: F = 51.09, df = 16, 357.5, p <.001; dog: 7.08 ± 0.45, human: 3.57 ± 0.45; contrast estimate = 3.50, t = 5.93, p <.0001; Figure 1 D), whereas food/water-borne transmission did not differ significantly between hosts (contrast estimate = 0.59, t = 1.01, p =.32). Dogs were also exclusive carriers of putative pathogens transmitted by inoculation (e.g. scratches, bites, such as *Pasteurella* sp.). Endogenous and food-transmitted pathogenic taxa were numerically more prevalent in human samples (38.7% ± 4% and 28.3% ± 3% respectively, Supp Figure 2C), though the endogenous route did not differ significantly between hosts in the mixed model.

**Figure 1.**
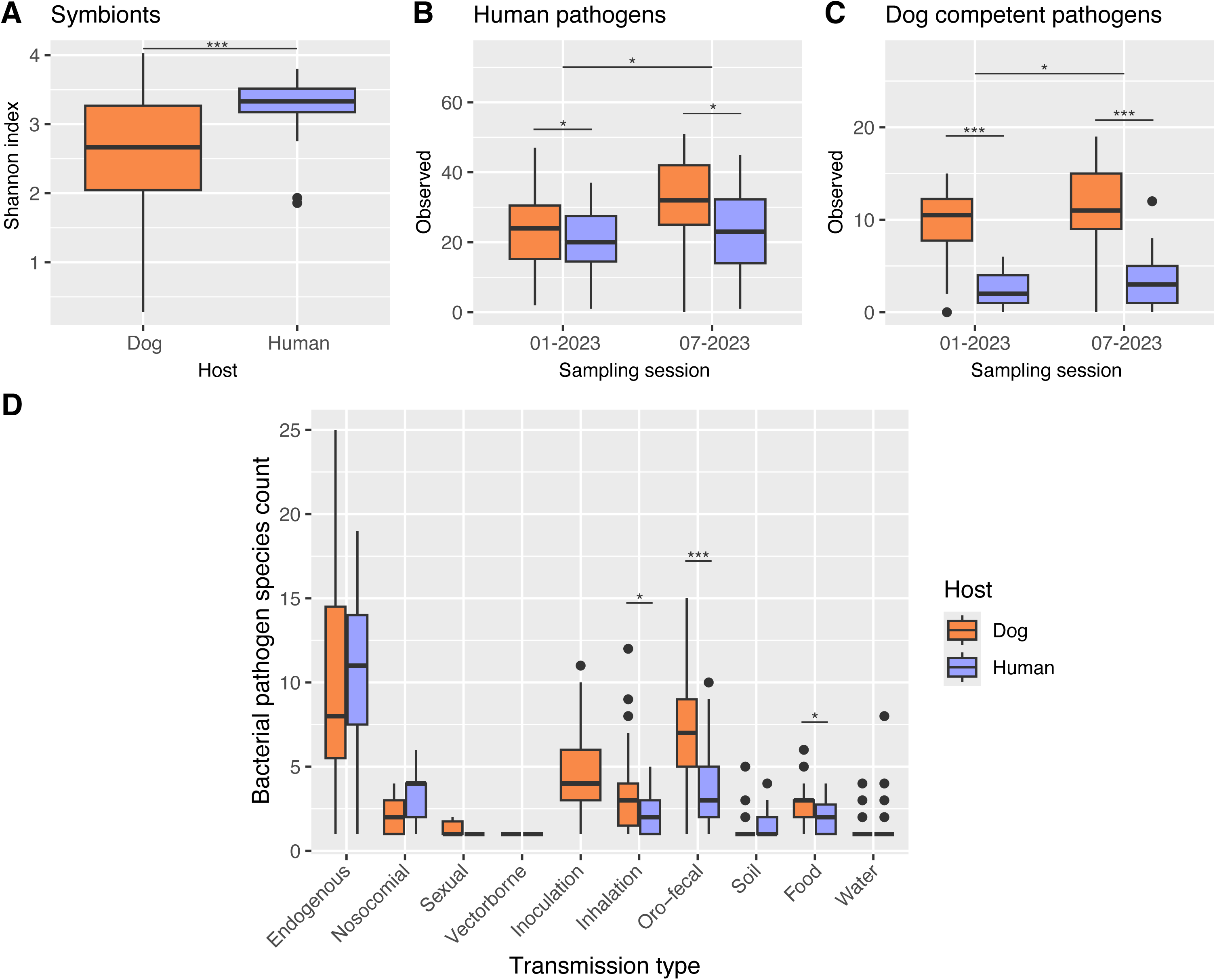
Alpha diversity boxplots of the (A) rectal symbionts, (B&D) human pathogens and (C) dog competent pathogens found in dog and humans of Nan; measured as the number of observed taxa at the lowest taxonomic level (B, C, D) and Shannon (A) index. The color code indicates whether samples originated from dogs (in orange) or humans (in purple), according to (B, C) sampling season or (D) transmission route. ∗,∗∗ and∗∗∗ represent significant contrasts results of restricted maximum likelihood fitting linear models p < 0.05, 0.01 and 0.001, respectively.

In general, dogs shared 16% of their symbionts with humans (n = 297 phylotypes), representing a great majority of their microbiome in terms of abundance (79% in number of reads). We observed very similar values for human pathogens, with 23% of shared taxa (n = 61), making up to 71 % of the number of reads in total for putative pathogens. When comparing the proportion of shared taxa between dog-owner pairs from within and outside their respective villages, we also observed similar results between gut symbionts and pathogens. For both types of bacteria, dogs share more taxa with other dogs than with humans, regardless of their village or respective owners (KW_symbionts_: χ2 = 731.58, df = 4, p <.001; KW_pathogens_: χ2 = 546.86, df = 4, p <.001; Figure 2). Dogs shared an average of 26.4% ± 1 % of their bacterial taxa with other dogs, and only 12.95% ± 4% with humans, without significant variations in shared taxa between dog-owner pairs and dogs paired with other humans from the same village or another location.

**Figure 2.**
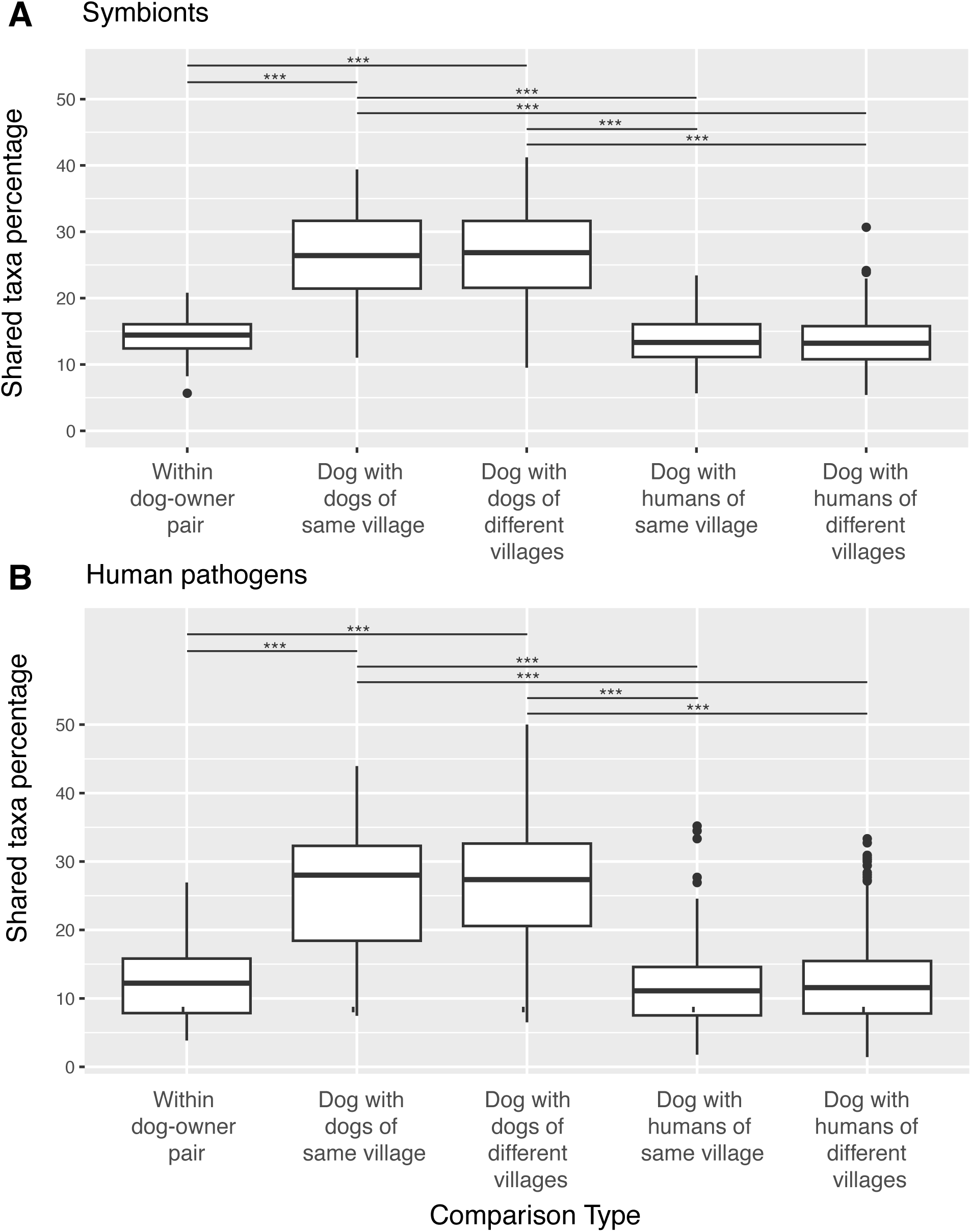
Boxplot of percentage of shared number of (A) symbionts taxa and (B) human pathogen species shared between dog-human pairs and dog pairs. ∗∗∗ = p < 0.001.

### 3.2 Microbial resemblance is highest among dogs, and stronger within dog-owner pairs than between dogs and other humans

Host type was a stronger driver of rectal microbiota structure compared to the season and residence: being either a dog or human explained for 30% and 24% of the variation in beta diversity among all samples for symbionts and putative pathogens respectively (PERMANOVA_symbionts_: F = 37.93, R2 =.304, df = 1, p <.001; PERMANOVA_pathogens_: F = 30.39, R2 =.24, df = 1, p <.001; Figure 3A) whereas their residence status explained less than 4% (PERMANOVA_symbionts_: F = 1.54, R2 =.037, df = 3, p <.001; PERMANOVA_pathogens_: F = 1.45, R2 =.034, df = 3, p <.001) and season less than 2% (PERMANOVA_symbionts_: F = 1.22, R2 =.01, df = 1, p =.232; PERMANOVA_pathogens_: F = 1.53, R2 =.012, df = 1, p =.12). The cause of the apparent clustering of human samples remains unclear.

**Figure 3.**
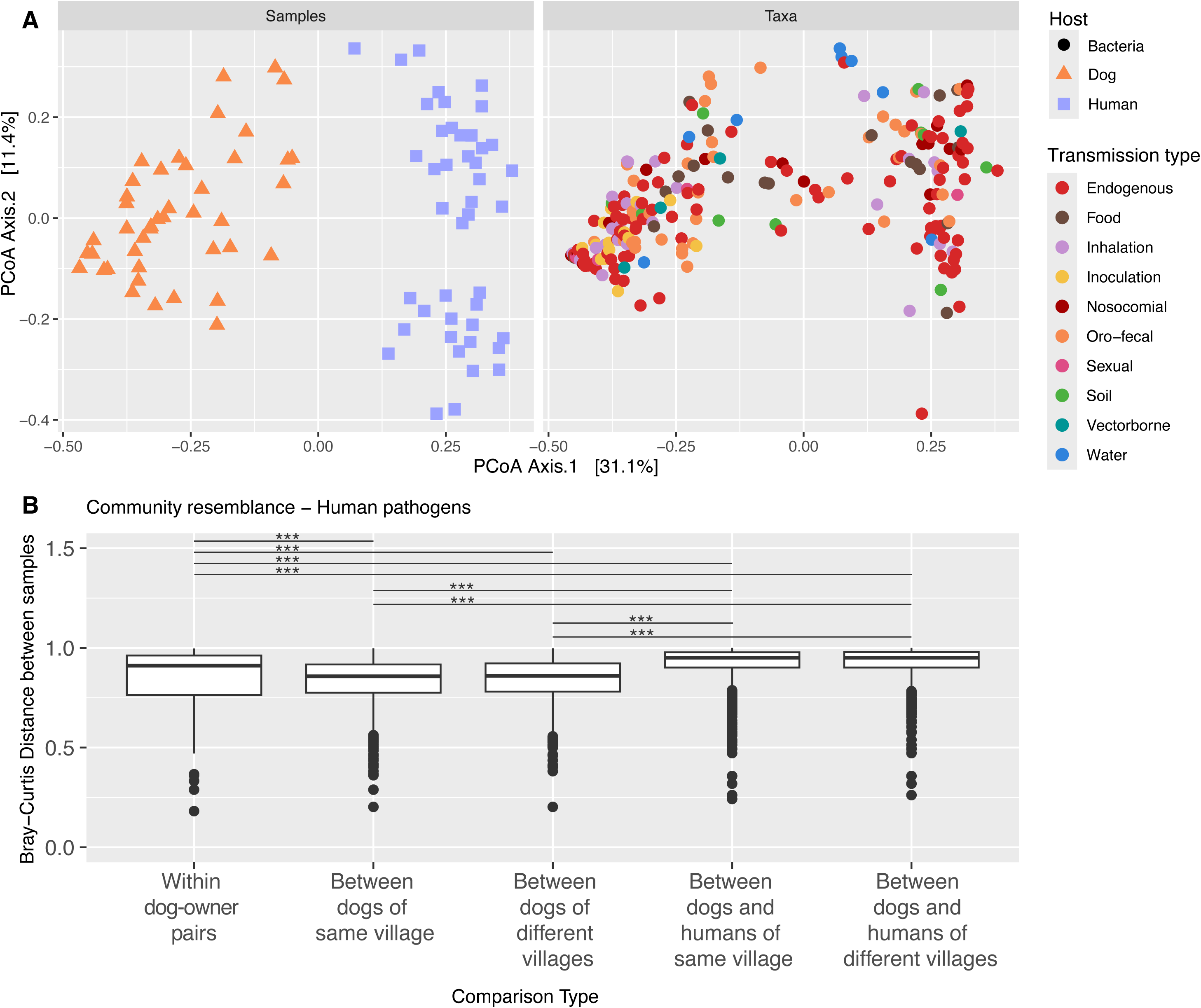
(A) PCoA biplot representing the beta diversity (calculated by Bray-Curtis distance) of the pathogenic bacteria from rectal microbiome. Left panel shows distribution of samples and right panel the distribution of bacterial taxa based on ordination. The color code corresponds to host type (left panel) and transmission route (right). (B) Boxplot of pathogen community resemblance based on Bray-Curtis distance between dog-human pairs and dog pairs. A distance of 0 represents exactly similar community between samples, 1 corresponds to absolutely different bacterial communities. ∗∗∗ represent p < 0.001.

As shown in the PCoA ordination based on Bray-Curtis distances between samples, pathogenic taxa transmitted through inoculation are more associated to dog samples (Figure 3A). Additionally, compositional variance was observed within dog and human samples for both types of bacteria (Betadisper_symbionts_: F = 14.48, df = 1, p <.001; Betadisper_pathogens_: F = 4.8, df = 1, p = 0.026). Variation in Bray-Curtis distances between samples also allowed us to investigate community resemblance between dog-human pair combinations. Within-species clustering (dog-dog) was stronger than cross-species in community resemblance, suggesting distinct host-specific microbiomes for both symbionts and putative pathogens (KW_symbionts_: χ2 = 1391.5, df = 4, p <.001; KW_pathogenss_: χ2 = 703.54, df = 4, p <.001). However, we observed greater community resemblance between dog-dog and dog-owner pairs than dogs with other humans regardless of their village of residence (Figure 3B).

### 3.3 Soil, food and oro-fecal transmitted taxa are more prevalent in dogs

According to our LDA effective size analysis, many putative pathogenic taxa were significantly different between dogs and humans: inoculation and soil-transmitted pathogens were more frequently found in dogs, while taxa transmitted through inhalation were associated with humans. Among the nine taxa differentially abundant in dogs and humans, seven of the nine taxa were transmitted through oro-fecal route and the two taxa transmitted through food were more abundant in dogs (Figure 4).

**Figure 4.**
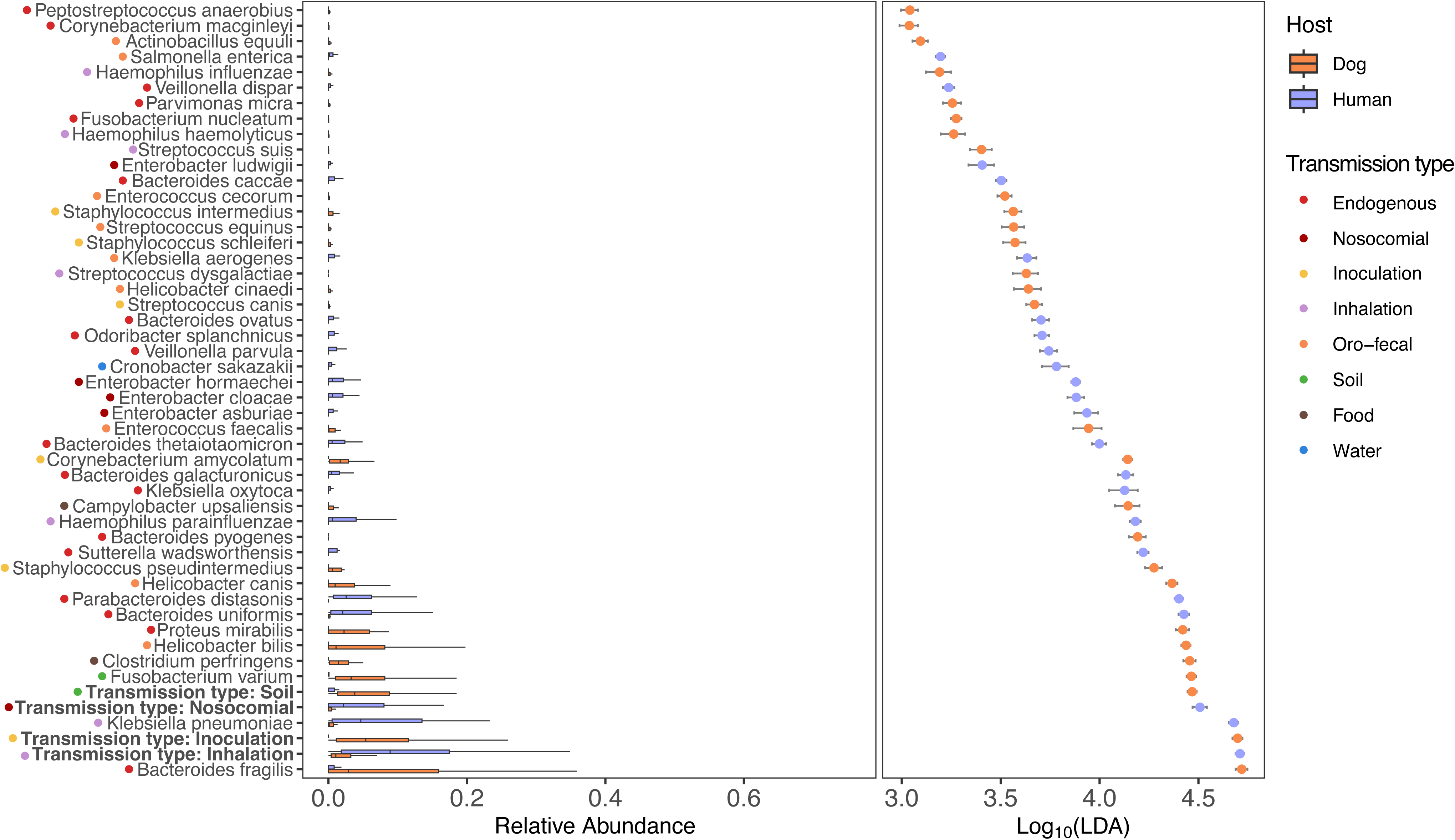
LDA scores (right) and boxplot of mean relative abundance (left panel) indicating statistical differences between microbial pathogens in dogs and humans. Data are presented from the species level and grouped by transmission routes (means of mean relative abundance by taxa). Colors indicate the host type with the highest mean of differential features in which significant differences were found.

Network analyses suggested differences in microbial association patterns between dogs and humans. For dogs the network structure indicated close associations within *Sutterella sp.* and *Prevotella sp.* taxa, as well as *Catenibacterium sp.,* while associations were suggested between *Enterococcus faecalis, Lactococcus garvieae, Lactococcus lactis* and *Aeromonas veronii*in humans (Figure 5). However, we observed high abundance of *E. coli* in both host types in association with *Shigella sp.* While overall resemblance was limited, a total of 131 taxa were common between dogs and their respective owners (when considering only taxa documented at least at the family level), including 98 symbionts, 24 known pathogens for humans and 8 zoonotic pathogens for which dog is competent. All were transmitted via food or oro-fecal route (*E. coli, Shigella sonnei, Shigella flexneri, Clostridium perfringens, Salmonella enterica, Shigella dysenteriae, Enterococcus faecium* and *Enterococcus faecalis*). We found co-occurrence of bacterial species within dog-owner pairs more than 50% of the time for 16 taxa only (Table 1), showing that dogs could potentially be proxies for their owners for *Fusobacterium mortiferum* (co-occurrence in pair for 73.3% of the cases) and *Klebsiella pneumoniae* (60.9%). On the other hand, putative pathogens with dogs as competent hosts had low paired co-occurrence (*S. sonnei* 33%, *S. flexneri* 31 %, *C. perfringens* 30%, *S. enterica* 20%, *S. dysenteriae* 10% *E. faecium* 7% and *E. faecalis* 6.7%). However, co-occurrence within dog-owner pairs cannot on its own distinguish whether dogs acquire these taxa directly from their owners, from a shared household environment, or independently from a common external source such as soil, water, or food; disentangling these pathways would require paired environmental sampling alongside dog and owner samples.

**Figure 5.**
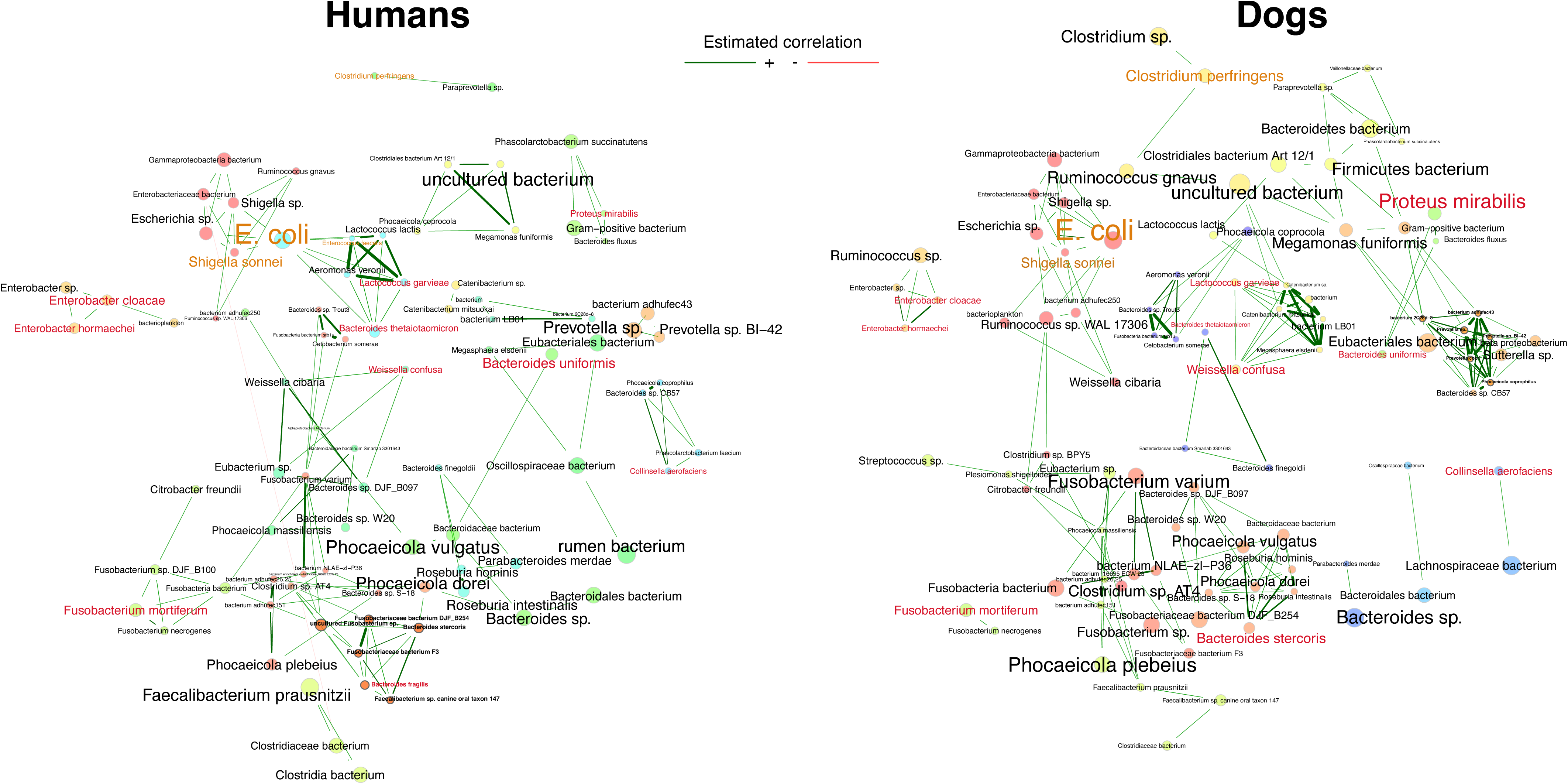
SPRING network associations within the microbiome of humans and dogs. Green edges represent positive associations, taxa names are sized based on overall abundance in all samples, red names correspond to human pathogens, and taxa in orange are taxa for which dogs are competent hosts. Node colors represent the different clusters identified.

**Table 1.** Occurrence of taxa found in common between dogs and their respective owners, and co-occurrence for each pair. Only taxa present more than 50% of the cases concomitantly in owners and their dogs.

| Bacterial species | Occurrence only in either dog or human within pair | Co-occurrence in dog-owner pair | Percentage of co-occurrence for pairs | Bacterial type |
| --- | --- | --- | --- | --- |
| *Escherichia coli* | 0 | 23 | 100,0 | Dog competent |
| *Fusobacterium mortiferum* | 6 | 17 | 73.3 | Human pathogen |
| *Klebsiella pneumoniae* | 9 | 14 | 60.9 | Human pathogen |
| *uncultured Ruminococcus sp.* | 2 | 19 | 90.5 | Symbiont |
| *uncultured Enterobacteriaceae bacterium* | 2 | 17 | 89.5 | Symbiont |
| *uncultured Clostridium sp.* | 5 | 17 | 77.3 | Symbiont |
| *uncultured Escherichia sp.* | 6 | 18 | 75,0 | Symbiont |
| *uncultured Fusobacterium sp.* | 6 | 16 | 72.7 | Symbiont |
| *uncultured Shigella sp.* | 7 | 15 | 68.2 | Symbiont |
| *uncultured Bacteroidaceae bacterium* | 8 | 15 | 65.2 | Symbiont |
| *uncultured Clostridiaceae bacterium* | 8 | 14 | 63.6 | Symbiont |
| *uncultured Haemophilus sp.* | 8 | 14 | 63.6 | Symbiont |
| *uncultured Eubacterium sp.* | 10 | 13 | 56.5 | Symbiont |
| *Ruminococcus gnavus* | 9 | 11 | 55,0 | Symbiont |
| *uncultured Streptococcus sp.* | 10 | 11 | 52.4 | Symbiont |
| *uncultured Sutterella sp.* | 10 | 11 | 52.4 | Symbiont |

## 4. Discussion

Several limitations should be noted. Our metabarcoding approach lacks strain-level resolution, likely overestimating pathogen prevalence since many species (e.g., *E. coli*) include commensal strains. The network analysis is exploratory and correlational: it visualizes potential co-occurrence patterns but does not test directionality, causality, or edge-level significance, so associations such as *E. coli*–*Shigella* co-occurrence should be treated as hypotheses for future targeted confirmation rather than established relationships. Our cross-sectional design (two time points, one year, 24 pairs, four villages) also limits our ability to generalize seasonal patterns or extend findings to other dog ecotypes and regions.

### 4.1 Microbial sharing and community resemblance

Overall, we found more diverse species in dogs’ microbiota compared to humans for both symbionts and pathogens, even though more evenness was found in human symbionts. Those results align with previous studies in dogs of multiple countries and lifestyles (Dunn et al., 2013; Kates et al., 2020; Misic et al., 2015), along with the remarkably high microbial diversity observed in Southeast Asian dogs (Yarlagadda et al., 2022). Seasonal changes were associated with variations in putative pathogen diversity, with dogs showing increased richness during the rainy season. These results align with ecological and climatic influences on microbial dynamics, potentially reflecting a heightened exposure to waterborne and soilborne taxa during wetter periods.

Compared to industrialized contexts, the free-ranging and “outdoor” lifestyles of dogs in Southeast Asia likewise amplify their exposure to diverse environmental microbes, contributing to these differences (Yarlagadda et al., 2022; Langlois et al., 2025). The shared taxa between dogs and humans highlight a potential zoonotic transmission risk, particularly for putative pathogens transmitted via oro-fecal routes. Dogs shared an average of 26.4% of their bacterial taxa with other dogs, but only 12.95% with humans regardless of their ownership and respective village, showing that the dog-owner relationship did not lead to substantial homogenization of the microbiota between the two host types. Proximity, shared environments, and behaviors such as grooming and cohabitation foster microbial transfer but also underscore host-specific factors that maintain distinct microbial communities. Indeed, a recent study on dog ownership practices in rural Thailand observed that most of the dogs had a free-ranging lifestyle, with 41.9% of respondents stating their dogs were always free-ranging and only 18.4% always restricting their dogs’ movements (Marshall et al., 2023). Free-roaming dogs are more likely to travel within and between villages to meet with other dogs.

We obtained similar results for community structures that were clearly distinct when comparing dogs to humans. However, individual dogs are closer to their owners than to other humans, aligning with the conclusions reached by most previous studies on the subject (Damborg et al., 2009; Ito et al., 2024; Song et al., 2013). This suggests that such patterns are not exclusive to westernized settings, as microbial similarities between dogs and their owners are also found in rural Thailand, distinguishing them from other individuals outside the household within the same village. However, this resemblance likely reflects shared household and community-level exposure — such as diet, water sources, and living space — rather than direct transfer of microbiota between a dog and its specific owner. Whether this pattern is primarily driven by shared dietary habits, particularly since most dog owners feed their pets leftovers and food scraps (Marshall et al., 2023), or by broader shared environmental exposure, remains to be determined.

### 4.2 Dogs as human proxies for zoonotic bacteria

While dogs harbored greater microbial diversity than humans and shared a subset of taxa, their utility as direct proxies for major local changes in human microbiota and outbreaks of pathogens appears limited. Instead, dogs may serve as indicators of shared exposure to specific zoonotic pathogens within a household or environment, rather than as direct proxies for their owner’s individual pathogen status. Potentially, high dog-dog interaction and exposure to diverse environments likely dilute the resemblance between dog and human microbiomes. However, the co-occurrence of putative zoonotic pathogens such as *Fusobacterium mortiferum* and *Klebsiella pneumoniae* in dog-owner pairs suggests that monitoring dog fecal bacteria could inform specific zoonotic disease dynamics, though this exploratory, community-level association would need confirmation through targeted, hypothesis-driven studies before being used for surveillance purposes. Indeed, the observed prevalence of *Shigella spp.* in both dogs and humans emphasizes the circulation of shared taxa linked to oro-fecal or food transmission route. This is particularly relevant for *Shigella spp.* for which humans are the primary reservoir (Percival & Williams, 2014), and thus dog-human contacts can explain high prevalence in dogs.

While targeted prevalence testing of true pathogenic taxa remains mandatory to confirm our results; these findings underscore the importance of integrating dogs into zoonotic disease surveillance and One Health initiatives, particularly in rural Southeast Asia. The dominance of putative pathogenic taxa transmitted via food and oro-fecal routes in dogs points to pathways for intervention. Enhanced public awareness and hygiene practices, alongside routine monitoring of companion animals, could help mitigate zoonotic risks. Moreover, this study highlights the necessity of accounting for cultural and environmental contexts when designing public health strategies, especially with human/animal interactions.

### 4.3 Future Directions

Longitudinal studies are needed to capture temporal variations in the microbiota and to elucidate causal relationships between environmental changes and microbial dynamics, as evidenced by our findings of increased potential pathogenic loads during the wet season. Expanding analyses to include targeted pathogen screening, with other dog ecotypes, different domestic animals and incorporating bacterial strain-level identification would offer deeper insights into pathogen-host interactions and enable more accurate risk assessments. Broadening the geographical and cultural scope of such research would further enhance our understanding of the factors driving microbial sharing and support the development of targeted interventions for the prevention of zoonotic diseases. Our study also reveals that dogs in rural Thailand exhibit microbial profiles consistent with the free-roaming dog ecotype, aligning with the dog-ecotype framework (Langlois et al., 2025).

In conclusion, while dogs may not fully mirror human microbiota, their microbiota offers a window into zoonotic risks and environmental microbial exposure. This study provides a foundation for future research exploring the intersection of human, animal, and environmental health in diverse contexts.

## Supporting information

Supp. mat.

## Acknowledgement

We would like to warmly thank for their cooperation the local authorities and the communities of Saenthong district, Nan province, Thailand.

## Data statement

The dataset has been deposited on CIRAD/Alfresco-SEAdogSEA project and may be accessed upon reasonable request to the data repository responsible person.

