## Supplementary material for "Exploring dog-owner microbiota and bacterial pathogens in Southeast Asia": Supp. mat.

**Supplementary Figure 1**. Rarefaction curve of all samples after mock and extraction blanks removal with red dashed line marking 20,000 reads.

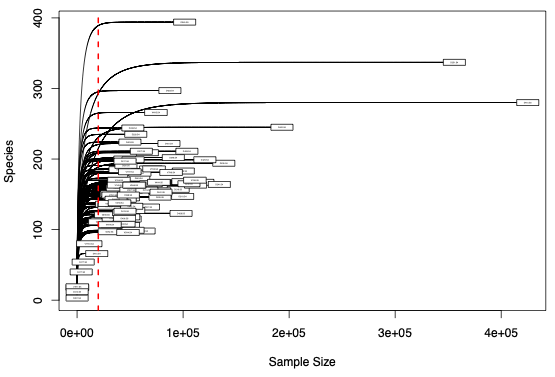

**Supplementary Figure 2**. Mean relative abundance of the most frequent bacterial phylotypes in the rectal microbiota of dogs-owner pairs, at the (A) phylum level per sampling period, (B) potential human pathogenic status and (C) potential pathogenic transmission route based on the existing literature. Samples are grouped according to their dog-owner pair based on their village of residence (H, Huak; HM, Huay Muang; N, NamKrai; S, Sun ti Suk).

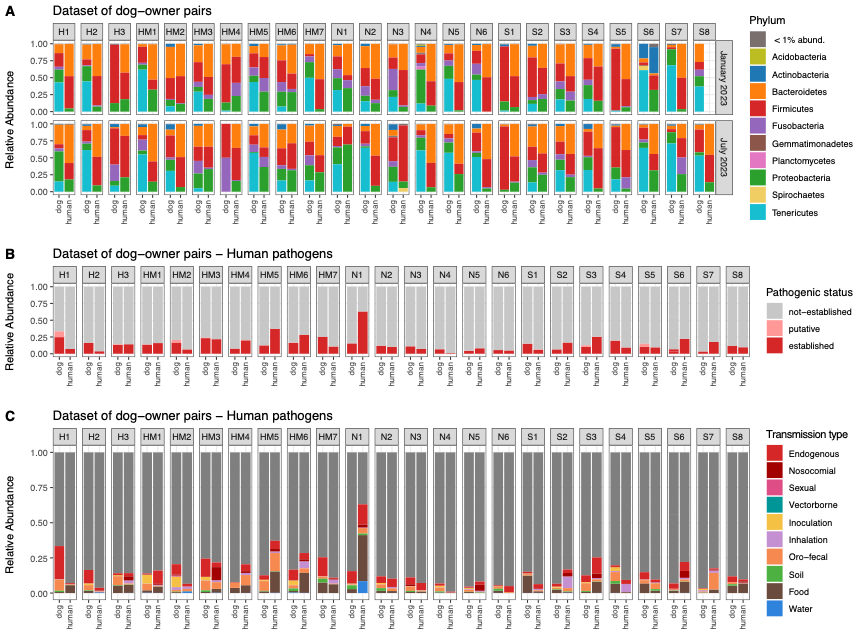

**Supplementary Table 1.** Sampling size per village according to dog-owner pairs. All of the dog-owner pairs were sampled twice (January and July 202

| **Village** | **Number of dog-owner pairs** | **Total number of samples** |  |
| --- | --- | --- | --- |
| Huak (4) | 3 | 12 |  |
| Huay Muang (6) | 7 | 27 | *One human sample removed* |
| Nam Krai (5) | 6 | 23 | *One dog sample removed* |
| Sun Ti Suk (7) | 7 | 25 | *One pair only sampled in july, other pair one dog sample removed* |
| **Total** | **23** | **87** |  |
